# Dynamic proton-dependent motors power Type IX secretion and gliding adhesin movement in *Flavobacterium*

**DOI:** 10.1101/2021.10.19.464928

**Authors:** Maxence S. Vincent, Caterina Comas Hervada, Corinne Sebban-Kreuzer, Hugo Le Guenno, Maïalène Chabalier, Artemis Kosta, Françoise Guerlesquin, Tâm Mignot, Mark McBride, Eric Cascales, Thierry Doan

**Author notes:** Current address: Department of Biochemistry, University of Oxford, South Parks Road, Oxford OX1 3QU, U.K.

## Abstract

Motile bacteria usually rely on external apparatus like flagella for swimming or pili for twitching. By contrast, gliding bacteria do not rely on obvious surface appendages to move on solid surfaces. *Flavobacterium johnsoniae* and other bacteria in the Bacteroidetes phylum use adhesins whose movement on the cell surface supports motility. In *F. johnsoniae*, secretion and helicoidal motion of the main adhesin SprB are intimately linked and depend on the type IX secretion system (T9SS). Both processes necessitate the proton motive force (PMF), which is thought to fuel a molecular motor that comprises the GldL and GldM cytoplasmic membrane proteins. Here we show that *F. johnsoniae* gliding motility is powered by the pH gradient component of the PMF. We further delineate the interaction network between the GldLM transmembrane helices (TMH) and show that conserved glutamate residues in GldL TMH are essential for gliding motility, although having distinct roles in SprB secretion and motion. We then demonstrate that the PMF and GldL trigger conformational changes in the GldM periplasmic domain. We finally show that multiple GldLM complexes are distributed in the membrane suggesting that a network of motors may be present to move SprB along a helical path on the cell surface. Altogether, our results provide evidence that GldL and GldM assemble dynamic membrane channels that use the proton gradient to power both T9SS-dependent secretion of SprB and its motion at the cell surface.

## Introduction

*Flavobacterium johnsoniae*, one of the fastest gliding bacteria described to date, uses surface-anchored adhesins to move on solid surfaces (Nelson, Bollampalli et al. 2008, Shrivastava, Rhodes et al. 2012, Nan, McBride et al. 2014). Remarkably, the major adhesin SprB exhibits a rotational behavior (Shrivastava, Lele et al. 2015) and its motion at the cell surface describes a closed helicoidal pattern along the long axis of the cell (Nakane, Sato et al. 2013, Shrivastava, Roland et al. 2016). It is proposed that binding of SprB to the substratum generates adhesion points and hence that SprB motion relative to the cell displaces the cell body in a forward screw-like motion (Shrivastava, Roland et al. 2016, Wadhwa and Berg 2021). SprB and other adhesins involved in gliding motility are transported to the cell surface by a multiprotein secretion apparatus, named type IX secretion system (T9SS) (Rhodes, Samarasam et al. 2010, Shrivastava, Johnston et al. 2013, Kulkarni, Johnston et al. 2019), which is present in most bacteria in the Bacteroidetes phylum (McBride and Zhu 2013, Abby, Cury et al. 2016).

The T9SS was discovered in the opportunistic pathogen *Porphyromonas gingivalis* in which it conveys a large number of virulence factors, including gingipain proteinases, across the outer membrane (OM) to the cell surface or the extracellular milieu (Sato, Naito et al. 2010, Goulas, Mizgalska et al. 2015, Nakayama 2015). In addition to gliding adhesins and gingipain proteinases, the repertoire of T9SS substrates also includes enzymes involved in nutrient supply and biofilm formation (Sato, Naito et al. 2010, Kharade and McBride 2014, Tomek, Neumann et al. 2014, Kita, Shibata et al. 2016). While the roles of the T9SSs and their substrates are relatively well known, information on T9SS architecture and mechanism of action are still sparse. However, conserved features of the T9SS have recently emerged (Lasica, Ksiazek et al. 2017, McBride 2019, Gorasia, Veith et al. 2020, Lunar Silva and Cascales 2021). The common T9SS architecture includes (i) the trans-envelope complex GldKLMN composed of two inner membrane (IM) proteins, GldL (or PorL) and GldM (or PorM), and of an OM-associated ring complex composed of the GldK OM lipoprotein and the GldN periplasmic protein (Gorasia, Veith et al. 2016, Vincent, Canestrari et al. 2017, Leone, Roche et al. 2018), (ii) the SprA (or Sov) OM translocon (Lauber, Deme et al. 2018), and (iii) the attachment complex that is comprised of the PorU, PorV and PorZ proteins (Chen, Peng et al. 2011, Glew, Veith et al. 2012, Gorasia, Veith et al. 2015, Glew, Veith et al. 2017, Madej, Nowakowska et al. 2021). These proteins assemble through a dense network of interactions that are poorly characterized and likely involve other conserved T9SS subunits.

T9SS-dependent secretion and gliding motility is a process energized by the IM proton-motive force (PMF) because inhibitors that dissipate the PMF prevent substrate secretion and halt cell displacement (Ridgway 1977, Pate and Chang 1979, Duxbury, Humphrey et al. 1980, Dzink-Fox, Leadbetter et al. 1997). At the single-cell level, Nakane and colleagues directly observed that SprB dynamics halted almost immediately after the addition of carbonyl cyanide *m*-chlorophenyl hydrazone (CCCP), a protonophore that collapses the PMF (Nakane, Sato et al. 2013). Hence, it was proposed that a PMF-dependent motor powers SprB dynamics and cell gliding. The nature of the molecular motor that powers SprB motion has been a longstanding question. Among the T9SS core components, only the GldL and GldM IM proteins share features with recognized PMF-dependent motors involved in the energization of flagellum rotation (MotAB), iron acquisition (ExbBD), outer membrane stability (TolQR), or myxococcal gliding motility and sporulation (AglQRS) (Block and Berg 1984, Skare and Postle 1991, Bradbeer 1993, Cascales, Lloubès et al. 2001, Sun, Wartel et al. 2011, Wartel, Ducret et al. 2013). GldL presents two transmembrane helices (TMHs) and largely faces the cytoplasm while GldM is a bitopic protein with a large periplasmic C-terminal domain. The *P. gingivalis* homologs of GldL and GldM (PorL and PorM, respectively) interact via their TMHs (Vincent, Canestrari et al. 2017). In addition, structural studies showed that the GldM and PorM periplasmic regions form dimers, and are composed of four domains, from D1 to D4 (Leone, Roche et al. 2018). The C-terminal D4 domain of PorM is involved in interactions with the outer membrane-associated PorKN complex (Leone, Roche et al. 2018). Finally, GldL/PorL and GldM/PorM bear conserved glutamate residues that may participate in harvesting the PMF (Vincent, Canestrari et al. 2017, McBride 2019). GldL and GldM are thus ideal candidates for constituting the IM proton-dependent motor powering type IX secretion and/or SprB dynamics. Indeed, a recent study presented the cryo-electron microscopy structure of the GldLM complex (Hennell James, Deme et al. 2021). The complex comprises two single transmembrane helices of GldM inside a pentameric ring of GldL, an architecture common with other known motors. The study also provided evidence that inter-TMH contacts modulated by the PMF are important for motor function. In addition, protonatable residues located in the GldL TMHs were shown to be essential for motor function (Hennell James, Deme et al. 2021). Here we provide further support and expand these conclusions. We show that the proton gradient component of the PMF is the source of energy powering gliding motility. We further demonstrate that the function of the GldLM motor requires a highly conserved glutamate residue in GldL, E49, whose protonation state controls interactions between the GldL and GldM TMHs and GldM conformation. We then show that substitution of a second GldL glutamate residue, E59, had no effect on secretion of SprB to the cell surface, but abolished SprB movement, thereby constituting a tool to uncouple T9SS-dependent secretion and gliding motility. Based on these results, we propose an updated model in which GldM conformational change upon PMF sensing is transmitted into mechanical torque through the periplasmic part of the T9SS to drive SprB motion.

## Results and Discussion

### Gliding is energized by the proton gradient

It is well known that gliding motility is arrested upon dissipation of the proton-motive force (Ridgway 1977, Pate and Chang 1979, Duxbury, Humphrey et al. 1980, Dzink-Fox, Leadbetter et al. 1997). The PMF consists of two gradients across the cytoplasmic membrane: an electrical potential (ΔΨ) and a chemical potential (ΔpH). To better define the energy source that powers gliding motility, *F. johnsoniae* gliding cells in a well chamber with glass bottom were subjected to valinomycin/K^+^ or nigericin, to specifically dissipate ΔΨ or ΔpH respectively, and single-cell gliding motility was quantified (Fig. 1A and B). In agreement with previous observations (Nakane, Sato et al. 2013), cells glide with an average speed of 1.7 µm.s^-1^ (Fig. 1B). As a control, injection of 10 µM of CCCP rapidly blocked all cell displacement in a reversible manner. By contrast, no significant inhibitory effect was observed upon addition of 40 µM valinomycin (+50 mM KCl). However, when cells were treated with 7 µM nigericin, motility was strongly impaired (Fig. 1B). Instead of gliding, cells appeared to jiggle around the same location, possibly because nigericin did not totally abolish the ΔpH (Fig. 1A). When nigericin was washed out, cells resumed gliding motility at normal speed. Therefore, we conclude that the proton gradient, but not the electrical potential, is the source of energy used by the gliding machinery.

**Figure 1.**
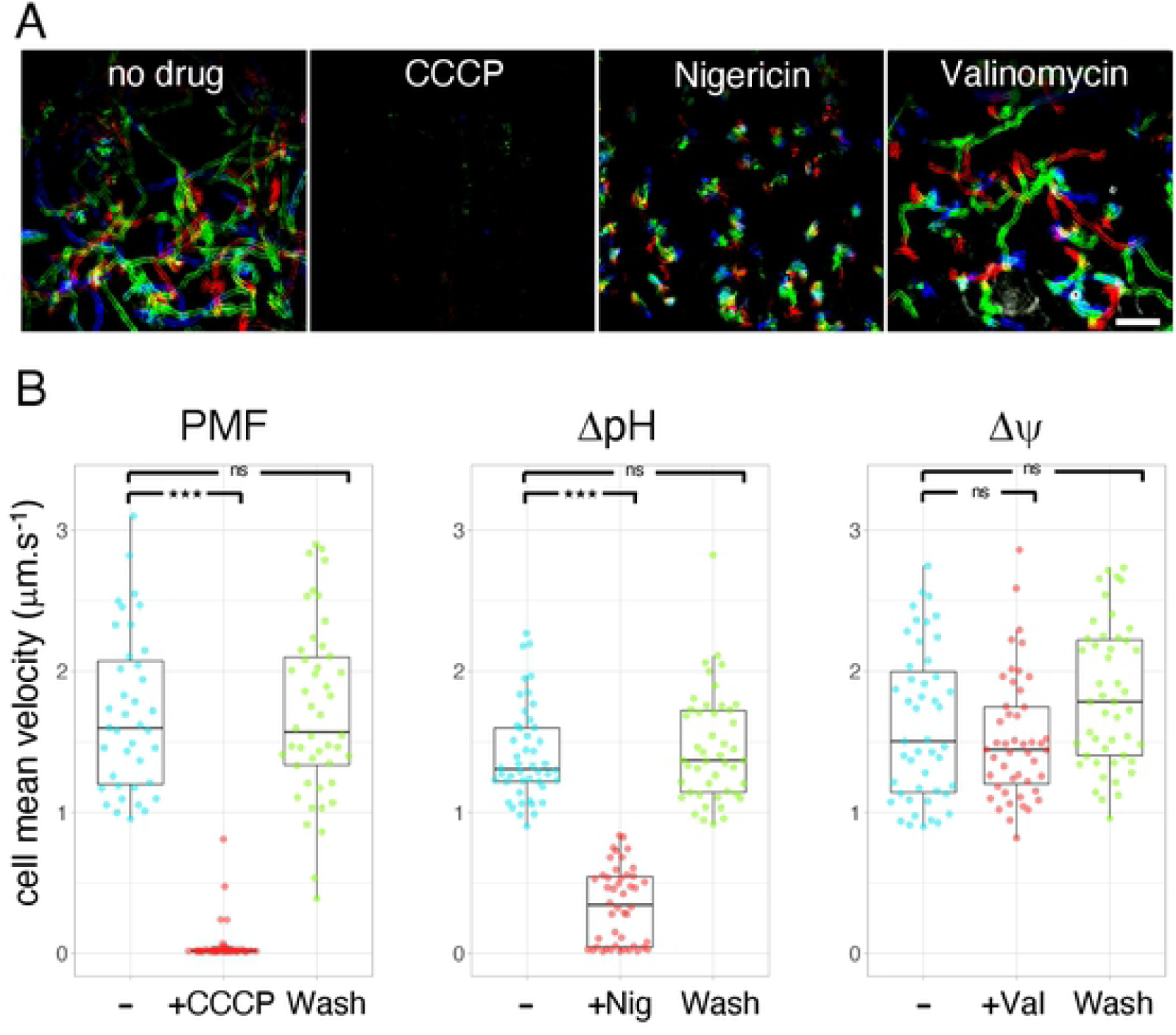
Effects of PMF dissipating drugs on the gliding of *F. johnsoniae* cells. **(A)** Rainbow traces of cell motility on glass recorded by phase contrast microscopy over time (2 min) in the absence of drug or in the presence of CCCP, nigericin or valinomycin. Individual frames from time lapse acquisition were coloured from red (start) to yellow, green, cyan and blue (end) and merged into a single rainbow image. Scale bar, 20 µm. **(B)** Combined jitter plots/boxplots of mean cell gliding velocity (in µm.s^-1^) of n>38 wild-type cells before (-), during a pulse of 10 µM CCCP (+CCCP), or 7 µM nigericin (+Nig), or 40 µM valinomycin/+50mM KCl (+Val), and after wash with fresh CYE medium (Wash). Statistical significance relative to the non-treated condition (-) is indicated above the plots (ns, non-significative; ^***^, *p* < 0.001; Wilcoxon’s *t*-test).

### GldL and GldM constitute the molecular motor that couples PMF to GldM conformational changes

Bacterial molecular motors such as the MotAB flagellar stator, the ExbBD and TolQR related transport systems and the AglRQS gliding motor generate mechanical energy by harvesting the chemical gradient through the cytoplasmic membrane (Blair and Berg 1990, Ahmer, Thomas et al. 1995, Cascales, Gavioli et al. 2000, Sun, Wartel et al. 2011). These complexes usually comprise two subunits organized in a 5:2 stoichiometry that interact via their TMHs (Celia, Botos et al. 2019, Santiveri, Roa-Eguiara et al. 2020). A conserved acidic residue, located in one TMH and facing the other TMHs, plays a key role in proton transit (Togashi, Yamaguchi et al. 1997, Zhou, Sharp et al. 1998, Celia, Noinaj et al. 2016, Celia, Botos et al. 2019, Santiveri, Roa-Eguiara et al. 2020). It is proposed that proton flow through the channel triggers protonation-deprotonation cycles of the side-chain of this residue and induces rearrangements in the TMHs, ultimately leading to the production of mechanical torque in the form of conformational changes in the extramembrane regions (Larsen, Thomas et al. 1999, Germon, Ray et al. 2001, Kojima and Blair 2001).

#### GldL and GldM interact via their transmembrane segments

In agreement with the recent cryo-EM structure of the GldLM complex, bacterial two-hybrid analyses show that GldL and GldM interact (Fig. 2A). As previously shown for the *P. gingivalis* PorLM complex (Vincent, Canestrari et al. 2017), these interactions likely involve the transmembrane segments of both proteins, as GldLM complex formation is prevented when only the soluble domains of these proteins are tested (Fig. 2A). Finally, similarly to the *P. gingivalis* PorKLMN complex, the GldLM module is implicated in interactions with the putative outer membrane-associated GldKN/O ring (Supplementary Fig. S1A) via contacts between the GldM periplasmic domain and GldK, GldN and GldO (Supplementary Fig. S1B). To test the contribution of TMHs to GldLM interactions, we conducted GALLEX and BLA approaches. GALLEX is based on the repression of a β-galactosidase reporter by two LexA DNA binding domains with different DNA binding specificities (LexA^WT^ and LexA^408^). If two TMHs interact, LexA^WT^ and LexA^408^ heterodimeric association causes repression of β-galactosidase synthesis (Schneider and Engelman 2003, Logger, Zoued et al. 2017). We tested interactions between TMHs displaying *in-*to-*out* topologies (GldL-TMH1 and GldM-TMH; Fig. 2B). GldL-TMH1 and GldM-TMH specifically formed homodimers but no interaction was detected between GldL-TMH1 and GldM TMH (Fig. 2C). To test the interaction with GldL-TMH2, which exhibits an *out-*to-*in* topology, we used the BLA assay (Julius, Laur et al. 2017, Schanzenbach, Schmidt et al. 2017). GldL-TMHs and GldM-TMH were fused to either the N- or C-terminal domain of β-lactamase (Bla). If an interaction between TMHs occurs, a functional β-lactamase is reconstituted and its activity can be quantified using a chromogenic substrate-based assay. In this assay, GldL-TMH2 homodimerization was not observed. However, GldL-TMH2 specifically interacted with both GldL-TMH1 and GldM-TMH (Fig. 2D). Taken together, our data show that GldL TMH1 and TMH2 interact with each other and GldL TMH2 interacts with GldM single TMH in the motor complex. With the exception of GldL-TMH1/GldL-TMH1 contacts detected by GALLEX, these data are in agreement with the position of the TMHs in the recent cryo-EM structure of the GldLM complex (Hennell James, Deme et al. 2021).

**Figure 2.**
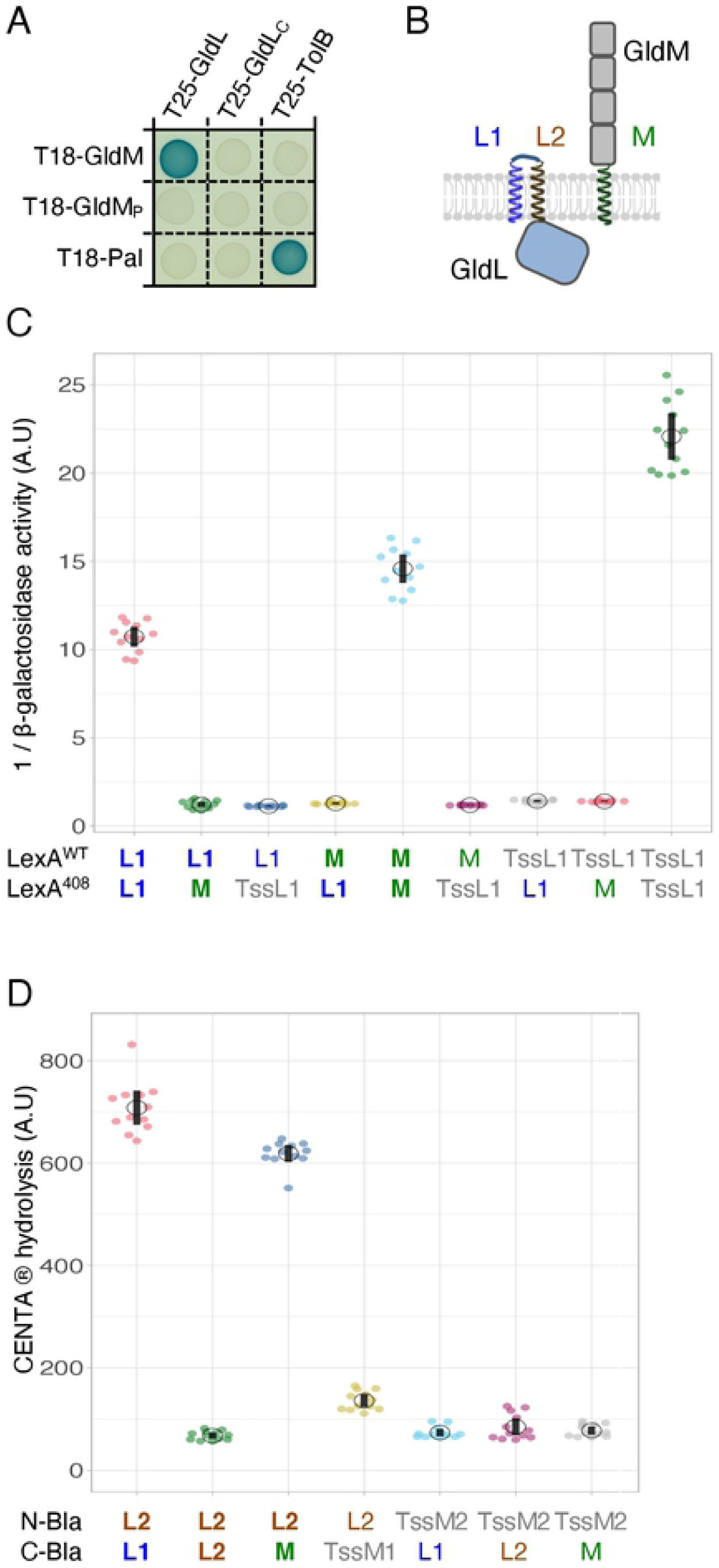
Interactions between GldL and GldM transmembrane helices. **(A)** Bacterial two-hybrid assay. GldL-GldM interaction is dependent on their TMHs. BTH101 cells producing the indicated proteins (GldL, GldM) or domains (GldL_C_, cytoplasmic domain of GldL, amino-acid 59 to 189; GldM_P_, periplasmic domain of GldM, amino-acid 36 to 513) fused to the T18 and T25 domain of the *Bordetella* adenylate cyclase were spotted on X-Gal-IPTG reporter LB agar plates. The blue color of the colony reports interaction between the two partners. Controls include T18 and T25 fusions to TolB and Pal, two proteins that interact but unrelated to the T9SS. **(B)** Schematic representation of GldL and GldM domains and topologies in the IM. The GldL TMH1 (L1, blue) and TMH2 (L2, brown), and GldM TMH (M, green) are indicated. **(C)** Homo- and heterodimerization of *in-*to*-out* TMHs of GldL and GldM probed with the GALLEX method. Jitter plots of β-galactosidase activity reporting the dimerization of TMHs fused to LexA^WT^ or LexA^408^. Measurements are reported as 1/ -galactosidase activity. Data are combined from technical triplicates of four independent measurements (2 colonies from two independent transformations each). Interactions with TssL1 (in grey) served as negative controls. TssL1 is the TMH of TssL, a protein of the *E. coli* Type VI secretion system (T6SS) that homodimerizes. **(D)** Homo and heterodimerization of the *in*-to-*out* and *out*-to-*in* THMs of GldL and GldM probed with the BLA method. Jitter plots of CENTA chromogenic substrate hydrolysis after 10 min of incubation. The activity is reported as the *A*_405nm_ value per *A*_600nm_. Controls include interaction assays with the TMHs of TssM (TssM1 and TssM2, in grey), a subunit of the *E. coli* T6SS.

#### GldM changes conformation depending on the proton gradient

We next sought to understand how the GldLM complex responds to the proton gradient. Our previous structural characterization of GldM and its homolog PorM from *P. gingivalis* revealed a conformational flexibility in the periplasmic region of GldM (Leone, Roche et al. 2018). The extracellular domain of GldM forms a straight homodimer that spans most of the periplasm. Each monomer is composed of four domains, D1 to D4 (Leone, Roche et al. 2018, Sato, Okada et al. 2020). Interestingly, the homolog PorM presents a kink between domains D2 and D3 (Leone, Roche et al. 2018), while a kink between the GldM D1 and D2 domains has been revealed in the cryo-EM structure (Hennell James, Deme et al. 2021), suggesting that GldM/PorM may alternate between several conformational states. Indeed, recent proteolytic susceptibility assays showed that the PMF regulates conformational changes in GldM: upon PMF dissipation by CCCP, two cleavages at the interface of domains D2 and D3 were identified by mass spectrometry after limited trypsinolysis (Song, Perpich et al. 2021). Here, we extend these observations by showing that the *in vivo* conformation of GldM is altered by drugs that perturb the proton gradient such as CCCP and nigericin, but remained unaffected upon treatment with the F_1_F_0_ ATPase inhibitors sodium azide and sodium arsenate, nor upon dissipation of the ΔΨ by valinomycin (Fig. 3A), demonstrating that GldM undergoes a structural transition dependent on the IM proton gradient (Fig. 3B).

**Figure 3.**
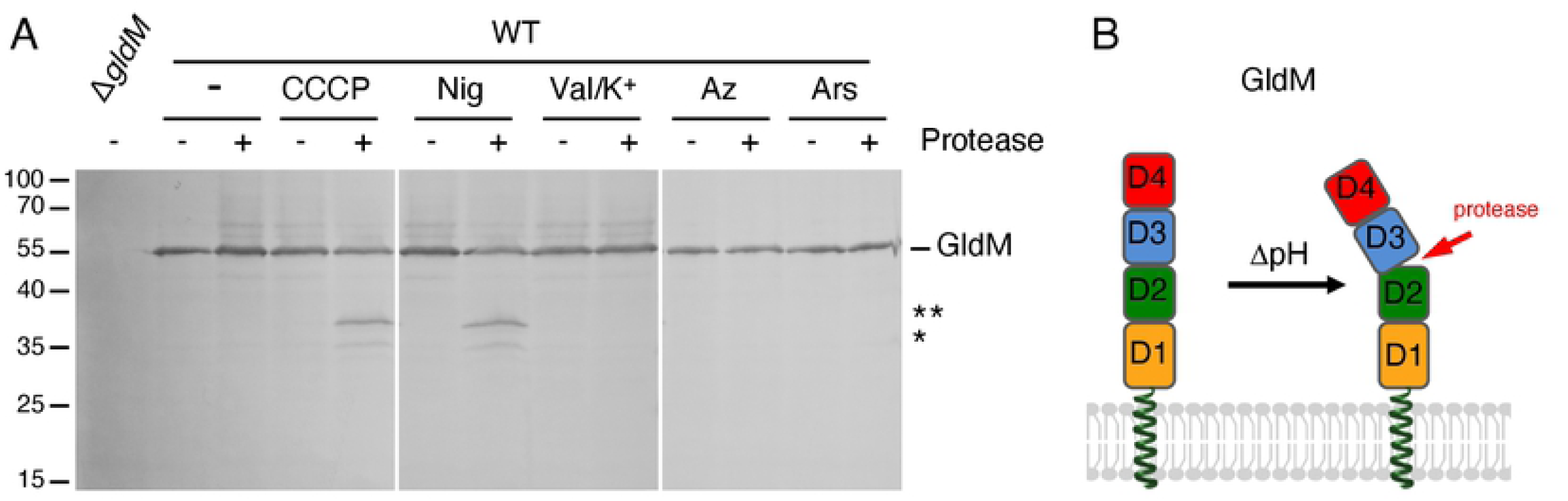
Conformational changes in GldM periplasmic domain in response to the proton gradient. **(A)** GldM protease accessibility assay. Spheroplasts of wild-type *F. johnsoniae* or the *gldM* mutant were treated (+) or not (-) with the trypsin protease and 10 µM CCCP (PMF inhibitor), 7 µM nigericin (pH inhibitor), 40 µM valinomycine/K+ (inhibitor), 1.5 mM sodium azide, or 20 mM sodium arsenate (F_1_F_0_ ATPase inhibitors). GldM was analyzed by SDS-PAGE and immunoblot with anti-GldM antibodies. The full-length GldM protein is indicated, as well as degradation products (^*^ and ^**^). The molecular weight markers (in kDa) are indicated on the left. **(B)** Schematic model of GldM conformational transition dependent on the proton gradient.

#### A conserved glutamate residue in GldL TMH2 is critical for harvesting the proton gradient

The activity of bacterial MotAB-like molecular motors characterized so far depends on conserved acidic residues located in their TMHs (Braun, Gaisser et al. 1996, Zhou, Sharp et al. 1998, Cascales, Lloubès et al. 2001, Sun, Wartel et al. 2011). We first tested the effect of the drug N-N’-Dicyclohexyl-carbodimide (DCCD), which covalently reacts with carboxylic groups located in a hydrophobic environment (Khorana 1953). Addition of DCCD abrogated gliding motility (Fig. 4A and B) and had the same effect on GldM conformation as CCCP (Fig. 4C). This effect was irreversible since DCCD remains covalently bound (Fig. 4B). These results therefore suggested that aspartate or glutamate residues are involved in coupling PMF to GldM conformational change and gliding motility. Sequence alignment showed that three acidic residues are conserved in the GldL and GldM N-terminal regions (Supplementary Fig. S2A and S2B): a glutamate at position 31 in the GldM TMH (GldM-E31) and two glutamates in GldL, one located in TMH2 (E49; strictly conserved in all GldL homologs in the OrthoDB database), and one located between TMH2 and the cytoplasmic domain (E59) (Supplementary Fig. S2C).

**Figure 4.**
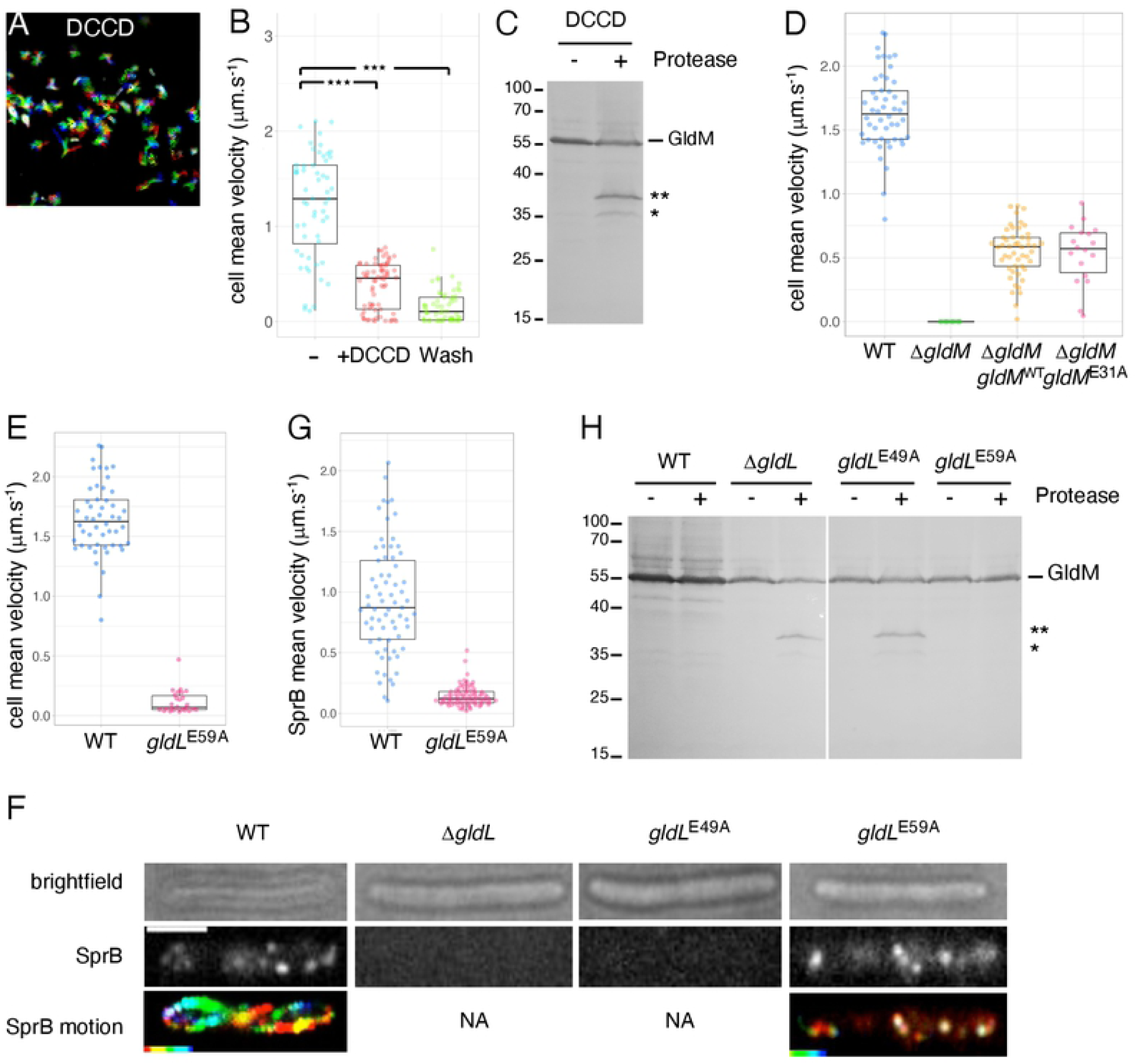
Roles of conserved GldL TMH2 glutamates in T9SS-dependent secretion and dynamics of SprB. **(A)** The addition of the drug DCCD inhibited cell motility. Rainbow traces of cell motility on glass recorded by phase contrast microscopy over time (2 min) in the presence of 10 µM DCCD. **(B)** Combined jitter plots/boxplots of mean cell gliding velocity (in µm.s^-1^) of n>50 wild-type cells before (-), during a pulse of 10 µM DCCD (+DCCD) and after wash with fresh CYE medium (Wash). Statistical significance relative to the non-treated condition (-) is indicated above the plots (ns, non-significative, ^***^, *p* < 0.001; Wilcoxon’s *t*-test). **(C)** Effect of DCCD on GldM protease susceptibility. GldM protease accessibility assay on spheroplasts of wild-type *F. johnsoniae* in absence (-) or presence (+) of 10 µM DCCD. **(D)** GldM conserved glutamate 31 does not play an important role in gliding motility. Combined jitter plots/boxplots of mean cell gliding velocity (in µm.s^-1^) of wild-type cells (WT, n=50), and strains expressing ectopically *gldM*^wt^ (n=49) or *gldM*^E31A^ point mutant (n=18) in a Δ*gldM* mutant background. The Δ*gldM* mutant has been placed in the graph for relevance but cell velocity has not been measured for that strain because it did not adhere to glass. **(E)** Combined jitter plots/boxplots of mean cell gliding velocity (in µm.s^-1^) of wild-type cells (WT, n=50) and a *gldL*^E59A^ point mutant (n=26). Strains were cultivated in CYE and single-cell gliding was observed on a free glass coverslip by phase contrast microscopy during 2 min. Gliding of the *gldL* mutant and of the *gldL*^E49A^ point mutant was not measured because cells did not adhere to the glass substratum. **(F**) Localization and dynamics of SprB on the cell surface in the wild-type strain (WT), a Δ*gldL* mutant, a *gldL*^E49A^ point mutant and a *gldL*^E59A^ point mutant. A representative cell is shown. Strains were cultured in CYE and, after SprB immunolabeling, were sandwiched between an agarose pad and a glass coverslip to significantly limit cell movement and facilitate SprB signal acquisition and analysis. SprB was immunolabeled using a primary serum directed against SprB and Alexa-488 fluorescent secondary antibodies. Fluorescence was recorded with 100 ms intervals for several seconds. The brightfield image (top panel), the first frame (middle panel, in grey levels) and the rainbow trace of SprB motion over time (bottom panel, not available for the Δ*gldL* mutant and the *gldL*^E49A^ point mutant) are shown. Scale bar, 2 µm. **(G)** Combined jitter plots/boxplots of mean displacement velocity (in µm.s^-1^) of SprB in wild-type cells (WT, n=69) and a *gldL*^E59A^ point mutant (n=85). SprB fluorescent spots were detected and tracked over time (>2 s) using the Trackmate plugin. **(H)** GldM protease accessibility assay in wild-type *F. johnsoniae* (WT), the Δ*gldL* mutant, and GldL^E49A^ and GldL^E59A^ point mutants. Spheroplasts were treated with (+) or not (-) with the trypsin protease. GldM was analyzed by SDS-PAGE and immunoblot with anti-GldM antibodies. The full-length GldM protein is indicated, as well as degradation products (* and ^**^). The molecular weight markers (in kDa) are indicated on the left.

Substitution of GldM E31 (GldM^E31A^) did not exhibit any defect in gliding motility compared to wild-type GldM, indicating that this residue does not play a significant role in T9SS-dependent secretion or gliding (Fig. 4D). By contrast, cells producing GldL^E49A^ failed to adhere to the glass surface, while substitution of GldL E59 abolished gliding motility without affecting adherence (i.e., SprB secretion by the T9SS) (Fig. 4E). Interestingly, the GldL^E49A^ and GldL^E59A^ variants presented distinct phenotypes regarding SprB secretion and dynamics as shown by live-cell immunolabelling using polyclonal anti-SprB antibodies and fluorescence time-lapse microscopy on agarose pads. As previously reported (Nakane, Sato et al. 2013), wild-type cells exhibited surface-exposed SprB fluorescent foci that describe an overall helicoidal pattern along the long cell axis, with dispersed velocity in the order of 1 μm.s^-1^ (Fig. 4F). No fluorescent focus was observed in Δ*gldL* mutant cells, which are unable to secrete SprB, indicating that SprB immunolabeling was specific. A similar observation was made with *gldL*^E49A^ mutant cells (Fig. 4F), demonstrating that T9SS-dependent SprB secretion to the cell surface requires residue E49 in GldL TMH2. By contrast, the GldL^E59A^ substitution supported SprB secretion but abolished the dynamic cell-surface movements of the adhesin (Fig. 4F-G). It is noteworthy that all GldL and GldM variants were produced in *F. johnsoniae* at levels comparable to the wild-type proteins (Supplementary Fig. S2D and S2E), although GldL^E49A^ migrated with lower apparent size than the wild-type protein (Supplementary Fig. S2E), an aberrant migration already observed in a separate study (Hennell James, Deme et al. 2021) and likely caused by the difference in detergent binding in SDS-PAGE between the TMH2 variants (Rath, Glibowicka et al. 2009). Taken together, these results support a model in which GldL E49 is required for secretion of the SprB adhesin and constitutes a key determinant of T9SS, whereas GldL E59 is dispensable for secretion and plays a specific function in gliding because it is only required for SprB movement. We next tested the contribution of these acidic residues for the regulation of GldM conformation. Protease accessibility assays showed that GldL and its Glu49 residue are required to maintain GldM in the conformation required for T9SS activity (Fig. 4H). By contrast, the GldL E59A substitution did not impact GldM proteolytic susceptibility (Fig. 4H), suggesting that the GldM conformation change observed by limited proteolysis is specifically linked to gliding motility rather than effector secretion.

#### Protonation of GldL glutamate residues

To address the question whether GldL E49 and E59 residues undergo protonation and deprotonation cycles, we determined their pKa values. A ^15^N/^13^C Glu-labeled synthetic peptide corresponding to GldL TMH2 (L2, residues Val40 to Val61) was solubilized in deuterated dodecylphosphorylcholine (DPC) micelles and analyzed by NMR. pKa values of 5.54±0.04 and 5.65±0.13 for the carboxylic groups of residues E49 and E59, respectively, were measured by the pH-dependent chemical shifts in two-dimensional ^13^C-HSQC experiments (Fig. 5A-B). In the presence of peptides corresponding to GldL TMH1 (L1, residues Lys6 to Thr29) and GldM TMH (M, residues Leu15 to Leu38), the pKa values slightly increased to 5.83±0.02 for both glutamates (Fig. 5B). The behavior of the ^13^C chemical shifts for the GldL TMH2 glutamate residues was then monitored in presence of the different peptides. At pH 5.2 (i.e., protonated glutamates), we observed chemical shift variations in the presence of L1, M or both (Fig. 5C). These data confirm that GldL TMH2 interacts with GldL-TMH1 and GldM-TMH, and that the presence of these TMH peptides influences the environment of the glutamate residues. However, at pH 6.7 (i.e., deprotonated glutamates), no chemical shift was observed upon addition of the L1, M or both peptides (Fig. 5C), suggesting that GldL TMH1 and GldM TMH are not in the environment of the glutamate residues. Taken together, these results suggest that the protonation state of the glutamic acids regulates contacts between TMH2 and the other TMHs in the GldLM complex, and hence that GldLM helix organization is likely to be modified during motor function, as evidenced for the MotAB and TolQR motors (Kim, Price-Carter et al. 2008, Zhang, Goemaere et al. 2009).

**Figure 5.**
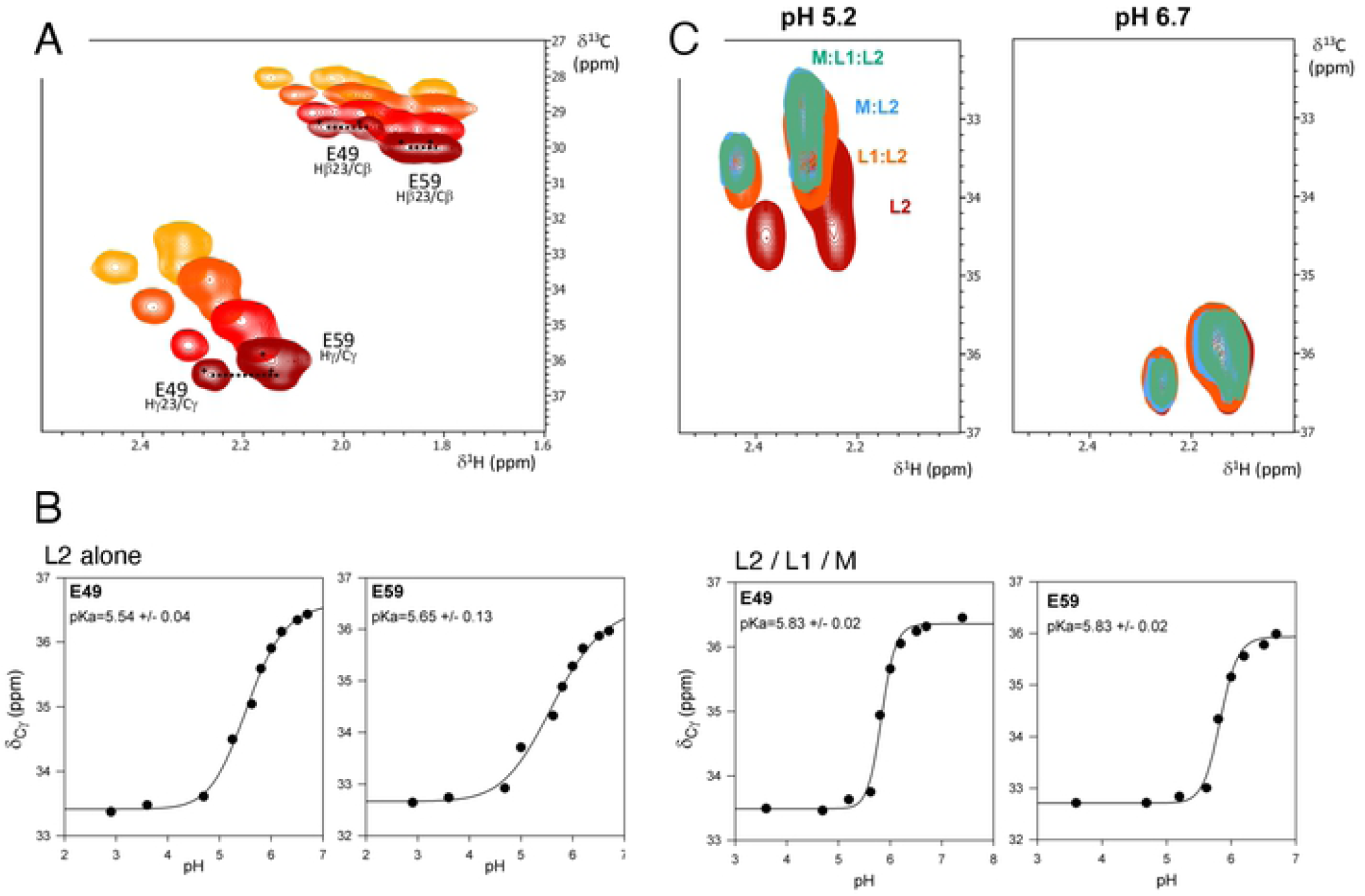
GldL glutamate residues protonation probed by NMR. pKa determination of ^13^C-Glu of free and complexed L2 peptide. **(A)** 2D ^13^C-HSQC spectra of 1 mM L2 peptide (^13^C-Glu labelled) in 150 mM deuterated DPC in 50 mM phosphate buffers at different pH (pH 2.9 (yellow), 5.0 (orange), 5.8 (red), 6.7 (brown)). **(B)** The pH dependent chemical shift variations of C γ carbons of E49 and E59 of the L2 peptide free, or complexed with L1 and M in a 1:1:1 molar ratio, were measured, fitted, and apparent pKa values were calculated using the Henderson-Hasselbach equation. **(C)** 2D ^13^C-HSQC spectra of 1 mM L2 peptide (^13^C-Glu labelled) in 150 mM deuterated DPC in 50 mM phosphate buffer at pH 5.2 (left panel) and pH 6.7 (right panel), in the absence (brown) and presence at molar ratio 1:1 of GldL-TMH1 peptide (L1, orange), GldM-TMH peptide (M, blue) and both L1 and M peptides (green).

Altogether our results support a model in which GldL and GldM form an IM proton channel with conserved critical glutamates that are protonated and deprotonated in response to the proton gradient to power both T9SS-dependent secretion and gliding motility. Our results also demonstrate that the protonation state of GldL E49 controls changes within the GldLM TMHs packing that are likely transmitted to the GldM periplasmic domain.

### GldLM motors and SprB adhesin do not have the same dynamics

SprB adhesins follow a closed right-handed helical track at the cell surface (Nakane, Sato et al. 2013, Shrivastava, Roland et al. 2016). Two models have been proposed on how the T9SS controls SprB motion (Nan, McBride et al. 2014, Shrivastava, Lele et al. 2015). In the first model, fixed rotary motors may activate treads to which SprB adhesins are connected. In the second model, SprB adhesins are directly connected to moving GldLM motors. The first scenario requires a network of motor complexes along the SprB helicoidal path. The second scenario implies that SprB colocalizes with dynamic GldLM complexes, and that SprB and GldLM move concomitantly along the helicoidal path. To explore these possibilities, we characterized the localization of the GldLM motor complex. Structured-illumination microscopy (SIM) recordings of *F. johnsoniae* fixed and permeabilized cells immunolabeled with polyclonal primary antibodies against the GldL, GldM, GldK or GldN proteins and fluorescent secondary antibodies showed that each protein was distributed in many foci along the cell body (Fig. 6A). For example, we numbered 32 GldL foci per cell in average (n=11, Fig. 6A). These data suggest that multiple Gld motors decorate the cell envelope. To provide further information, we sought to perform live observations. However, none of the plasmid-borne or chromosomal fluorescent fusions to GldL or GldM we generated supported wild-type gliding, possibly due to the size of the fluorescent protein tags. We therefore turned to a more sophisticated method to generate functional and time-trackable proteins, using the alfa technology (Götzke, Kilisch et al. 2019).

**Figure 6.**
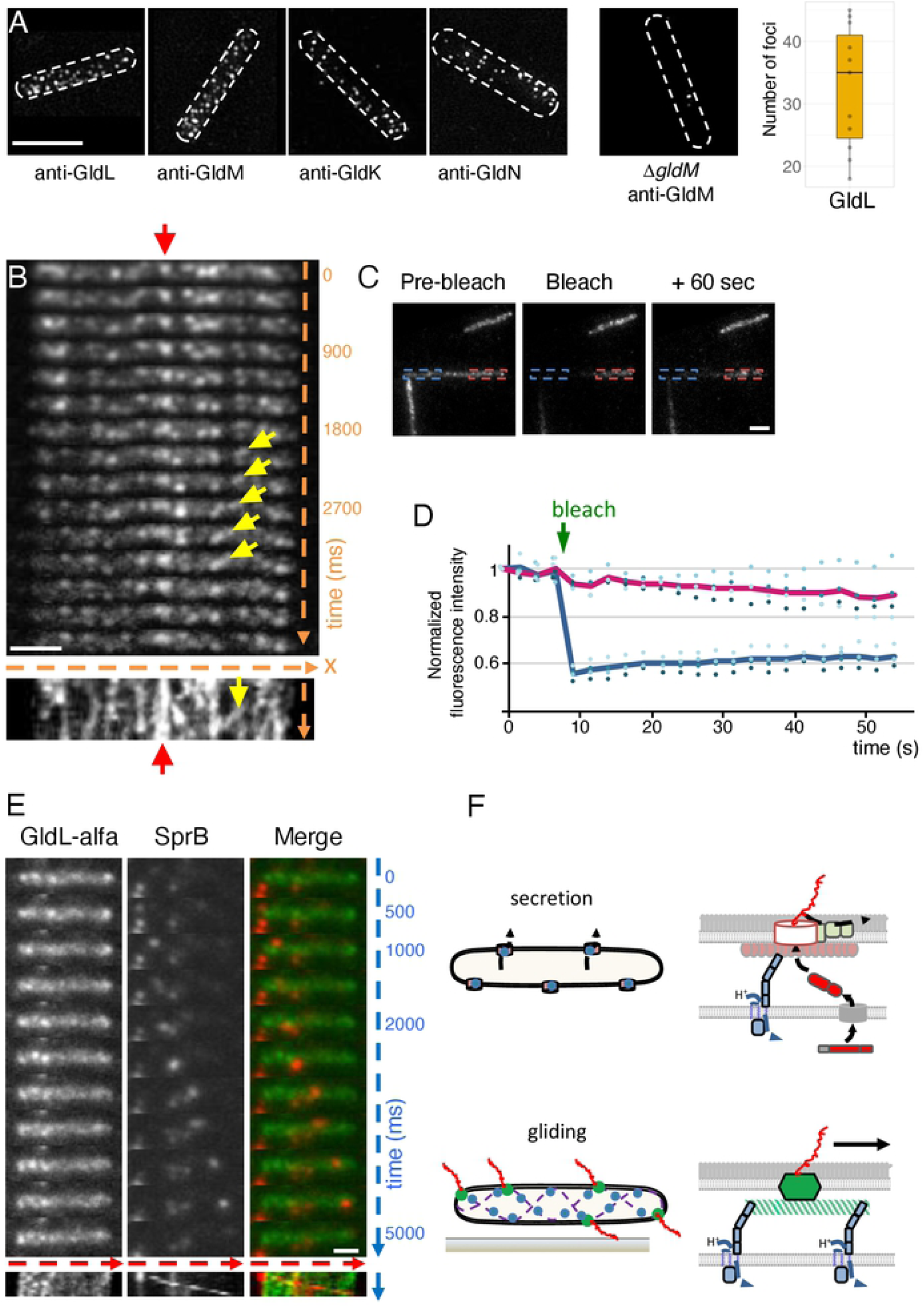
Dynamic localization of Gld complexes. **(A)** Localization of Gld proteins in fixed cells. Immunostaining of GldL, GldM, GldK and GldN and observation by structured-illumination microscopy. A typical specificity control is shown on the right panel with a Δ*gldM* mutant stained with antibodies directed against GldM. Scale bar 2 µm. The number of foci was quantified for GldL-alfa and shown in a boxplot. **(B)** Live-cell dynamics of a functional GldL-alfa fusion bound to NbAlfa-sfGFP. After NbAlfa-sfGFP induction, GldL-alfa dynamics was followed by time lapse fluorescence microscopy with 300 ms intervals. For a representative cell, a stack of individual frames is shown. Time is indicated in ms. At the bottom of the stack, a kymograph of the fluorescence signal in the same representative cell. The x-axis is the position of GldL-alfa/NbAlfa-sfGFP signal with respect to the substratum (glass), and the y-axis is time. The red arrow shows an example of a static signal and the yellow arrow points to an example of a moving GldL-alfa focus. Scale bar 2 µm. **(C)** GldL-alfa does not travel long distance within the cell. Diffusion of GldL-alfa/NbAlfa-sfGFP fluorescence signal over time assayed by FRAP. Cells expressing GldL-alfa and NbAlfa-sfGFP were pulse-bleached in the region indicated by the blue rectangle. A representative cell is shown. Scale bar 2 µm. **(D)** Fluorescence intensities were measured in the bleached region (blue rectangle) and a non-bleached region (red rectangle) for 60 sec in 4 cells. Individual measurements are shown with spots (filled spots for bleached regions and empty spots for non-bleached regions). The green arrow indicates the moment of bleaching. The fluorescence intensity of each region of interest was normalized to the first prebleached intensity. The blue (bleached region) and red (non-bleached) lines indicate the average of all measurements. **(E)** GldL-alfa and SprB do not colocalize. Dynamic localization of GldL-alfa bound to NbAlfa-sfGFP (in green) and immunostained SprB (in red) in a representative cell. Fluorescence was followed by hilo microscopy with 500 ms intervals. A stack of individual frames is shown. Time is indicated in ms. At the bottom of the stack, a kymograph of the fluorescence signal in the same representative cell. The x-axis is the position of the cell with respect to the substratum (glass), and the y-axis is time. Scale bar, 1 µm. **(F)** Model of GldLM molecular motors function in Type IX secretion (top) and surface adhesin dynamics (bottom). GldLM motors (blue) are fueled by the proton gradient, leading to conformational shifts of the periplasmic domain of GldM. When associated to the T9SS (top), GldLM motors generate mechanical torque to rotate a GldKNO ring to drive secretion. GldLM motors may also be associated to the gliding machinery (bottom), in which they serve to transport SprB (in red) on the cell surface via the displacement of a track or baseplate machinery (in green).

The alfatag is a 13-amino-acid peptide that is specifically and almost irreversibly bound by the NBalfa nanobody with an affinity of ∼0.26 pM (Götzke, Kilisch et al. 2019). The sequence encoding the alfatag was introduced in frame with the GldL-coding sequence at the native locus. GldL-alfa was functional and supported single-cell gliding (Fig. S3A-B). We then engineered a replicative plasmid expressing NBalfa-sfGFP under the control of an IPTG-inducible promoter in *F. johnsoniae*. Expression of NBalfa-sfGFP did not perturb cell gliding, either in a wild-type background or in a strain expressing GldL-alfa (Supplementary Fig. S3B). NBalfa-sfGFP was diffuse in wild-type cells that do not express GldL-alfa (Fig. S3C). By contrast, NBalfa-sfGFP exhibited a punctate pattern in GldL-alfa cells (Fig. 6B). Distinct foci were visible as well as more patchy signals, rendering quantification difficult. Remarkably, these foci were not all static relative to the cell, as opposed to the SprA translocon (Lauber, Deme et al. 2018), nor did they behave like SprB adhesins that travel along the entire cell length. Time-lapse microscopy and kymograph analyses of signal dynamics indicated that some foci remain static while others moved quickly but at varying speed relative to the cell (Fig. 6B and Movie S1). In addition, fluorescence recovery after photobleaching (FRAP) experiments in non-moving cells suggest that GldL-alfa movement was restricted to short distances within the cell because fluorescent signal could not be recovered over a large bleached cell region (Fig. 6C-D). These two GldL populations could correspond to GldLM complexes engaged into static complexes with the OM translocon to secrete substrates, and to free GldL proteins or to GldLM complexes following a track to energize SprB motion. However, colocalization experiments in live cells with immunolabeled SprB showed that GldL-alfa and SprB do not follow the same trajectories (Fig. 6E). These results support the tread model (Nan and Zusman 2016, Shrivastava and Berg 2020) in which GldLM proton channels convert the proton gradient into mechanical force to displace or activate treads involved in SprB movement (Fig. 6F).

### Concluding remarks

In this study, we provided evidence that Type IX Secretion and surface adhesin motion are energized by a molecular motor fueled by the proton gradient, like the flagellar motor and other bacterial molecular motors. Our data support the idea that interactions between the transmembrane helices of GldL and GldM shift in response to the proton flux, eventually leading to conformation changes in the GldM periplasmic domain. Conserved glutamate residues in GldL are important in this process but are not equivalent in terms of function. While amino acid E49 in GldL is essential for secretion through the T9SS as observed by Hennell and colleagues (Hennell James, Deme et al. 2021), glutamate at position 59 is only required for gliding, indicating that in *F. johnsoniae*, T9SS secretion itself does not require concomitant SprB motion along the cell surface. Thus, it is tempting to speculate that secretion and SprB motion are not supported by the same mechanical rearrangements in GldM, or that SprB motion may require more mechanical torque than the secretion process. Furthermore, since SprB motion and secretion are uncoupled in the GldL E59A point mutant, this mutation is an interesting tool to study secretion independently of gliding in *F. johnsoniae*. Nevertheless, GldL-E59 is also conserved in non-gliding bacteria like *P. gingivalis*, suggesting that it may also serve for T9 secretion in other bacteria (Supplementary Fig. S2A).

Our data also support the idea that GldM conformational shift upon PMF sensing could be converted into mechanical torque through the periplasmic part of the T9SS. Indeed, we showed that GldM periplasmic domain is connected to the T9SS GldKNO subcomplex, similar to the PorKLMN complex in *P. gingivalis*. Two recent studies help understand how this could work. First, the structure of the GldLM motor showed that ten GldL TMHs (five GldL molecules) wrap two GldM TMHs in an asymmetric manner (Hennell James, Deme et al. 2021). It was proposed that GldM TMHs would rotate within a GldL ring in response to the PMF to generate mechanical movement of GldM periplasmic domain. These findings are consistent with our data and provide an explanation for why GldL-E49 is required for motor function. However, they do not explain the role of E59, which is located outside the membrane in the GldLM structure (Supplementary Fig. S2C; Hennell James et al., 2021). One may hypothesize that E59 enters the proton channel when GldM rotates. Second, *in situ* PorKN rings were observed by cryo-electron tomography (Gorasia, Chreifi et al. 2020, Song, Perpich et al. 2021). These rings may serve to maintain T9SS subcomplexes in close proximity to allow sequential translocation, maturation and attachment of the substrates (Gorasia, Chreifi et al. 2020). Therefore, an attractive hypothesis is that GldM conformational changes in response to the proton gradient could generate mechanical torque for the rotation of GldKNO rings, similar to cogwheels, that directly or indirectly facilitate secretion of T9SS substrates.

Finally, our results are consistent with the “rack and pinion” model proposed by Shrivastava and Berg to explain how the GldLM complex participates in SprB displacement (Shrivastava and Berg 2020). Our microscopy data suggest the existence of static GldLM motors, which are presumably associated with static T9SS translocons, and GldLM complexes that are dynamic but that move differently than do SprB molecules. These motors could be linked to unidentified motion treads carrying SprB adhesins.

## Material and Methods

### Bacterial strains, media and chemicals

All strains are listed in Table S1. *Escherichia coli* strains DH5α and BTH101 were used for cloning procedures and bacterial two-hybrid assay, respectively. *E. coli* cells were grown in Lysogeny Broth, at 37°C or 28°C. For BACTH experiments, gene expression was induced by the addition of iso-propyl-β-D-thio-galactopyranoside (IPTG, Sigma-Aldrich, 0.5 mM) and plates were supplemented with 5-bromo-4-chloro-3-indolyl-β-D-galactopyranoside (X-Gal, Eurobio, 40 µg.mL^-1^). *F. johnsoniae* CJ1827, a streptomycin-resistant *rpsL*2 derivative of ATCC 17061 (UW101), was used as model micro-organism. *F. johnsoniae* cells were grown at 28°C in Casitone Yeast Extract (CYE) medium (Agarwal, Hunnicutt et al. 1997) or Motility Medium (MM) (Liu, McBride et al. 2007) as indicated. For selection and maintenance of the antibiotic resistance, antibiotics were added to the media at the following concentrations: erythromycin, 100 µg.mL^-1^; streptomycin, 100 µg.mL^-1^; tetracycline, 20 µg.mL^-1^, ampicillin, 100 µg.mL^-1^, kanamycin, 50 µg.mL^-1^, chloramphenicol, 40 µg.mL^-1^. Specific enzyme and chemicals source were as follows: trypsin (Sigma), Carbonyl cyanide *m*-chlorophenyl hydrazone (CCCP, Sigma, 10 µM), nigericin (Nig, Sigma, 7 µM), valinomycine (Val, Sigma, 40 µM), sodium azide (Az, Sigma, 1.5 mM), arsenate (Ars, Sigma, 20 mM), *N,N*′-Dicyclohexylcarbodiimide (DCCD, Sigma, 100 µM).

### Genetic constructs

All plasmids and oligonucleotide primers used in this study are listed in Table S1. Enzymes for PCR and cloning were used as suggested by manufacturers.

Chromosomal mutants were generated as described (Rhodes, Pucker et al. 2011). The suicide plasmid designed to generate an in-frame deletion of *gldM* was built as follows. A 2.5 kb fragment containing the region upstream of *gldM* and *gldM* start codon was PCR amplified using oligonucleotide primers F1-ΔgldM and R1-ΔgldM. This fragment was digested with *Bam*HI and *Sal*I and inserted into plasmid pRR51 cut with the same restriction enzymes to generate an intermediate plasmid. A 2.5 kb fragment containing *gldM* stop codon and the region downstream of *gldM* was PCR amplified using oligonucleotide primers F2-ΔgldM and R2-ΔgldM. Similarly, it was digested with *Sal*I and *Sph*I and inserted into the previously generated plasmid cut with *Sal*I and *Sph*I to generate plasmid pRR51-Δ*gldM*.

The suicide plasmids designed to build *gldL*^E49A^ and *gldL*^E59A^ strains were constructed as follows. A plasmid with plasmid pRR51 backbone and carrying a 4 kb region centered around *gldL* E49A and E59A codon substitutions was synthesized (Geneart, Thermofisher). The suicide plasmid pRR51-*gldL*^E49A^ was then built by restoring the codon for E59 in the previously synthesized plasmid by site directed mutagenesis with oligonucleotide primers Fw-GldL-A59E and Rv-GldL-A59E. Similarly, the suicide plasmid pRR51-*gldL*^E59A^ was built by restoring the codon for E49 in the previously synthesized plasmid by site directed mutagenesis with oligonucleotide primers Fw-GldL-A49E and Rv-GldL-A49E.

The suicide plasmid used to generate the GldL-alfa fusion was made as follows. A 1.5 kb fragment containing *gldL* and part of the alfa tag was PCR amplified using oligonucleotide primers oTM582 and oTM590. A 1.5 kb fragment containing part of the alfa tag, a stop codon and the region immediately downstream of *gldL* stop codon was PCR amplified using oligonucleotide primers oTM591 and oTM592. These fragments were assembled using Gibson isothermal reaction and reamplified using oligonucleotide primers oTM582 and oTM592. This fragment was digested with *Bam*HI and *Sph*I and inserted into pRR51 cut with the same restriction enzymes. Plasmids for GALLEX and BLA were engineered by hybridizing complementary oligonucleotides corresponding to the GldL or GldM TMHs, and inserting them into *Nhe*I-*Bam*HI-digested target GALLEX or BLA vectors. BACTH plasmids were engineered by restriction and ligation as previously described (Vincent, Canestrari et al. 2017). The replicative plasmid designed for complementation and production of GldLM^WT^ were constructed as follows. A fragment containing *gldL* and *gldM* open reading frames was PCR amplified using oligonucleotide primers 5-BamHI-LM and 3-XbaI-LM. This fragment was digested with *Bam*HI and *Xba*I and inserted into plasmid pCP11 (McBride and Kempf 1996) cut with the same restriction enzymes. The replicative plasmid designed to produce GldM^E31A^ was then generated by quick change site directed mutagenesis using oligonucleotide primers 5-GldM-E31A and 3-GLdM-E31A.

The replicative plasmid designed to express NbAlfa-sfGFP from an IPTG inducible promoter in *F. johnsoniae* was designed as follows. A first replicative plasmid was built with an IPTG-inducible promoter, a multicloning site and *lacI* constitutive expression for repression in the absence of IPTG in *F. johnsoniae*. A fragment containing the promoter of Fjoh_0697 (Chen, Kaufman et al. 2010) with *lacO3* and *lacO1* operator sites flanking the −33 and −7 promoter sequences, pCP23 multicloning site and Fjoh_0139 promoter after the *Pst*I restriction site was synthesized (Geneart, Thermofisher). A fragment containing *lacI* open reading frame was PCR amplified using oligonucleotide primers oTM495 and oTM496. These fragments were assembled by Gibson isothermal reaction and reamplified using oligonucleotide primers oTM497 and oTM496. This fragment was digested with *Kpn*I and *Sph*I and inserted into pCP23 (Agarwal, Hunnicutt et al. 1997) cut with the same restriction enzymes to generate plasmid pCP-*lac*. Then, the gene encoding NbAlfa was synthesized (Geneart, Thermofisher) and reamplified using oligonucleotide primers oTM612 and oTM596. sfGFP, codon-optimized for translation in F. johnsoniae, was also synthesized (Geneart, Thermofisher) and then reamplified by PCR using oligonucleotide primers oTM595 and oTM593. These fragments were assembled using the Gibson isothermal reaction and reamplified using oligonucleotide primers oTM612 and oTM593. It was then digested with *Bam*HI and *Nhe*I and inserted into plasmid pCP-*lac* cut with the same restriction enzymes.

### Protein interaction assays

The adenylate cyclase-based bacterial two-hybrid technique was used as previously published (Vincent, Canestrari et al. 2017). Briefly, the proteins to be tested were fused to the isolated T18 and T25 catalytic domains of the *Bordetella* adenylate cyclase. After introduction of the two plasmids producing the fusion proteins into the BTH101 reporter strain, plates were incubated at 28°C for 24 h. Three independent colonies for each transformation were inoculated into 600 µL of LB medium supplemented with ampicillin, kanamycin, and IPTG (0.5 mM). After overnight growth at 28°C, 10 µL of each culture was spotted onto LB plates supplemented with ampicillin, kanamycin, IPTG, and X-Gal and incubated at 28°C. Controls include interaction assays with TolB and Pal, two protein partners unrelated to the T9SS. The experiments were done in triplicate and a representative result is shown.

GALLEX and BLA were performed as described (Logger, Zoued et al. 2017).

### Protease susceptibility assay

*F. johnsoniae* cells were grown in 5 mL of CYE medium to an *A*_600_=0.8, harvested by centrifugation and resuspended in 100 µL of 20 mM Tris-HCl pH 8.0, 20% sucrose, 1 mM EDTA, and 100 µg.mL^-1^ of lysozyme. After 30 min incubation at room temperature (20°C), 100 µL of ice-cold sterile water was added and the mixture was carefully mixed by three inversions. 50 µL of each spheroplast suspension was treated with trypsin (100 µg.mL^-1^). After 5 min on ice, 17 µL of boiling 4× Laemmli loading buffer was added, and immediately vortexed and boiled for 5 min prior to SDS-PAGE and immunoblot.

### Western blot analyses

*F. johnsoniae* cells were grown to mid-log phase in CYE at 28°C. Whole cells were prepared for SDS-PAGE and Western blotting assays were performed as previously described. Equal amounts of total proteins were loaded for each sample based on culture optical densities. Anti-GldL, anti-GldM (Shrivastava, Johnston et al. 2013) and anti-FLAG (Sigma Aldrich, clone M2) antisera were used at 1/5000, 1/5000 and 1/10,000 dilutions, respectively.

### Nuclear Magnetic Resonance

NMR experiments were carried out on a Bruker Avance III 600 MHz spectrometer, at 300 K. Three synthetic peptides L1 (GldL-TMH1: KKVMNFAYGMGAAVVIVGALFKITKK), L2 (GldL-TMH2: KKVMLSIGLLT**E**ALIFALSAF**E**PVKK) and M (GldM-TMH: KKLMYLVFIAMLAMNVSK**E**VISAFGLKK), with ^15^N/^13^C-Glu-labeled, have been studied free and in complexes at molar ratio 1:1 or 1:1:1. NMR samples containing 1 mM peptide concentration in 150 mM deuterated DPC were used in different phosphate buffers (50 mM). The behaviour of the ^13^C chemical shifts for glutamate residues in the different peptides as a function of pH (2.9 to 8.9) was monitored using a two-dimensional ^13^C-HSQC experiment. Chemical shift values as a function of pH were analyzed according to a single titration curve of the form:

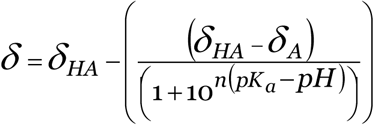

where *δ* is the observed chemical shift at a given pH, *δ*_HA_ and *δ*_A_ are the chemical shifts for the various protonated forms of the peptide, and *n* is the number of protons transferred.

### Fluorescence microscopy and image analysis

#### General microscopy

For single-cell gliding on glass, cells were grown in CYE at 28°C to an *A*_600 nm_ ≈ 0.7. Cells were diluted to an *A*_600 nm_≈ 0.05 and 100 µL were spotted into µ-Slide chambers with glass coverslip bottom (Ibidi). After 5-min incubation, floating cells were washed out with fresh CYE medium and gliding of adherent cells was monitored by phase contrast microscopy on a Nikon Eclipse TE-2000 microscope equipped with a 100× NA 1.3 Ph3 objective, a perfect focus system to maintain the plane in focus, and an Orcaflash 4.0 LT digital camera (Hamamatsu Photonics). GldL-alfa/NBalfa-sfGFP localization was observed by Hilo microscopy. Cells were grown in CYE overnight without shaking at 28°C. NBalfa-sfGFP expression was induced with 1 mM IPTG for 1 h prior to observation. Cells were spotted on a 2 % low-melting agarose pad for immediate observation. Hilo fluorescence microscopy and FRAP experiments were performed with a Nikon Eclipse Ti2 microscope equipped with a 100x NA 1.45 Ph3 objective, an Orca-Fusion digital camera (Hamamatsu), a perfect focus system, and an Ilas2 TIRF/FRAP module (Gataca Systems).

#### Immunolabelling and SIM acquisition

SprB immunolabelling on live cells was performed essentially as described (Nakane, Sato et al. 2013). Briefly, 500 µL of cells were incubated 5 min with a 1/100 dilution of antiserum directed against SprB (Nelson, Bollampalli et al. 2008). Cells were washed once with CYE and further incubated 5 min with Alexa488- or Alexa561-labeled anti-rabbit secondary antibodies (Thermofisher). Cells were washed four times in CYE and concentrated 5-fold. In order to facilitate SprB detection and tracking during short periods, cells were spotted on a 2 % low-melting agarose pad for immediate observation. Immunolabeling of GldL, GldM, GldK and GldN was performed on fixed cells as previously described (Braun and McBride 2005), except cells were manipulated in tubes instead of on glass slides. Polyclonal antisera directed against GldL, GldM, GldK or GldN (Shrivastava, Johnston et al. 2013) were used at 1/2000 dilution and further recognized by Alexa488-labeled anti-rabbit secondary antibodies (Thermofisher). Structure illumination microscopy (SIM) was performed on a DeltaVision OMX SR microscope (GE Healthcare). The experiments were done in triplicate and a representative result is shown.

#### Image analysis

Images were analyzed using ImageJ (http://imagej.nih.gov/ij/). The MicrobeJ plugin (Ducret, Quardokus et al. 2016) was used to detect and track cells during gliding. The Trackmate plugin (Tinevez, Perry et al. 2017) was used to detect SprB fluorescence and analyze its dynamics. Statistical dataset analysis was performed using Excel and the R software environment (https://www.r-project.org/). Kymographs were generated using the KymoResliceWide plugin (https://imagej.net/KymoResliceWide, E. Katrukha and L. Young).

For fluorescence recovery quantification, images were corrected for bleaching using histogram matching prior to signal recovery quantification.

## Data availability

All data and material are made available upon request.

## Acknowledgements

We thank the members of the Cascales team for insightful discussions and support, Jean-Pierre Duneau for discussion regarding peptide solubilization, Moly Ba, Isabelle Bringer, Annick Brun, Olivier Uderso, Mathilde Valade and Audrey Gozzi for technical assistance. This work was supported by the Aix-Marseille Université (AMU), the Centre National de la Recherche Scientifique (CNRS), and grants from the Agence Nationale de la Recherche (ANR-15-CE11-0039 and ANR-20-CE11-0017), from the Excellence Initiative of Aix-Marseille University (A*MIDEX, A-M-AAP-ID-17-33-170301-07.22), a French “Investissements d’Avenir” programme, and from the Fondation Bettencourt-Schueller. M.S.V. was supported by a doctoral fellowship from the French Ministère de la Recherche, and an end-of-thesis fellowship from the Fondation pour la Recherche Médicale (FDT2018-05005242).

## Author Contibutions

MSV, EC and TD conceived the study. MSV, CCH, CSK, HLG, MC, EC and TD contributed to experiments and data analysis, with the main contribution of MSV. AK, FG, TM and MM provided resources, technical and conceptual input. EC acquired funding. MSV, EC and TD wrote the original draft. CSK, MSV, EC and TD edited the manuscript with the help of other co-authors.

## Legend to Supplementary Figures

**Supplementary Figure S1** | **Network of interactions between proteins of the T9SS core components**. Bacterial two-hybrid assays. **(A)** T9SS outer membrane-associated core complex (GldK, GldN and GldO) and GldJ. **(B)** T9SS outer membrane-associated core complex (GldK, GldN, GldO), GldJ and the inner membrane-associated core complex (GldM, GldL). The signal sequence was omitted in the constructs for GldN and GldO. The signal sequence and the acylated N-terminal cysteine residue of the mature form were omitted for GldK and GldJ. BTH101 reporter cells producing the indicated proteins or domains (GldL_C_, cytoplasmic domain of the GldL protein; GldM_P_, periplasmic domain of the GldM protein) fused to the T18 or T25 domain of the *Bordetella* adenylate cyclase were spotted on plates supplemented with IPTG and the chromogenic substrate X-Gal. The TolB-Pal interaction serves as positive control. **(C)** Model of the interactions between T9SS components defined by bacterial two-hybrid.

**Supplementary Figure S2** | **(A)** Sequence alignments of the N-terminal regions that encompass the two transmembrane segments of GldL homologs. The alignment was performed using TCOFFEE. Red arrows indicate the conserved acidic residues. **(B)** Sequence alignments of the region that encompasses the single transmembrane segment of GldM homologs. The alignment was performed using TCOFFEE. The red arrow indicates the conserved acidic residue. **(C)** Highlight of GldL-E49 (orange) and E59 (pink) glutamate residues in the structural model of the GldLM complex. The left panel shows a side view and the right panel shows a view from the cytoplasm. GldL TMHs are colored green. GldM subunits (TMH and first periplasmic domain) are colored blue. **(D)** Western blot analysis of GldM production using anti-GldM antibodies in a Δ*gldM* mutant, wild-type *F. johnsoniae* (WT), GldM WT or GldM E31A expressed from a plasmid in a Δ*gldM* mutant background. **(E)** Western blot analysis of GldL production using anti-GldL antibodies in the Δ*gldL* mutant (Δ*gldL*), wild-type *F. johnsoniae* (WT), and strains expressing GldL^WT^-flag (GldL^WT^) or GldL^E49A^-flag (E49A) or GldL^E59A^-flag (E59A). Extracts of cells were subjected to SDS-PAGE and immunodetection with anti-GldL and anti-Flag primary antibodies and HRP-coupled secondary antibodies. Molecular weight markers (in kDa) are indicated on left.

**Supplementary Figure S3** | **GldL-alfa supports cell gliding. (A)** Rainbow traces of cell motility on glass recorded by phase contrast microscopy over time (2 min) in a wild-type strain and a strain expressing *gldL-alfa* at the native locus. Individual frames from time lapse acquisition were coloured from red (start) to yellow, green, cyan and blue (end) and merged into a single rainbow image. Scale bar, 20 µm. **(B)** Combined jitter plots/boxplots of mean cell gliding velocity (in µm.s^-1^) of *gldL-alfa* cells in the absence of NbAlfa-sfGFP (-, n=124) or with 1 mM IPTG induction of NbAlfa-sfGFP (+, n=125) or wild-type cells in the absence of NbAlfa-sfGFP (-, n=135) or with 1 mM IPTG induction of NbAlfa-sfGFP (+, n=151). **(C)** Representative micrograph of cells expressing fluorescent NbAlfa-sfGFP in a wild-type background. Scale bar, 2 µm.

